# Thyroid Hormone Suppresses Medulloblastoma Progression Through Promoting Terminal Differentiation of Tumor Cells

**DOI:** 10.1101/2024.02.13.580111

**Authors:** Yijun Yang, Silvia Anahi Valdés-Rives, Qing Liu, Yuzhe Li, Jun Tan, Yinfei Tan, Christian A Koch, Yuan Rong, Steven R. Houser, Shuanzeng Wei, Kathy Q Cai, Sheue-yann Cheng, Tom Curran, Robert Wechsler-Reya, Zeng-jie Yang

**Author notes:** To whom correspondence may be addressed: Zeng-jie Yang.

## Abstract

Hypothyroidism is commonly detected in patients with medulloblastoma (MB). A possible link between thyroid hormone (TH) signaling and MB pathogenicity has not been reported. Here, we find that TH plays a critical role in promoting tumor cell differentiation. Reduction in TH levels frees the TH receptor, TRα1, to bind to EZH2 and repress expression of NeuroD1, a transcription factor that drives tumor cell differentiation. Increased TH reverses EZH2-mediated repression of NeuroD1 by abrogating the binding of EZH2 and TRα1, thereby stimulating tumor cell differentiation and reducing MB growth. Importantly, TH-induced differentiation of tumor cells is not restricted by the molecular subgroup of MB. These findings establish an unprecedented association between TH signaling and MB pathogenicity, providing solid evidence for TH as a promising modality for MB treatment.

## Introduction

Medulloblastoma (MB), the most prevalent malignant brain tumor among children, comprises at least four molecular subgroups: Wingless (WNT), Sonic Hedgehog (SHH), Group 3 (G3) and Group 4 (G4). Each subgroup exhibits unique clinical and genetic characteristics ^1,2^. The SHH subgroup predominantly affects infants and young adults and is marked by germline or somatic mutations in genes associated with Hedgehog (Hh) pathway signaling ^3^, such as loss-of-function mutations or deletions in *Patched 1* (*Ptch1*) or activating mutations in *Smoothened* (*Smo*). G3 is the most aggressive MB subgroup, with a 5-year overall survival rate of less than 60% ^4^. Despite comprehensive multimodal therapy, a significant proportion of patients continues to succumb to this disease, and survivors often suffer enduring treatment-related side effects including endocrine disorders and cognitive deficits ^5,6^. Consequently, there exists a pressing need for more effective and less toxic approaches to treat MB.

Previous studies have illuminated a key aspect of MB biology: MB tumor cells can undergo terminal differentiation, a process marked by an irreversible exit from the cell cycle and the loss of tumorigenic potential ^6-8^. This suggests that inducing terminal differentiation in MB tumor cells may offer a viable therapeutic strategy. However, to harness this potential, it is imperative to fully understand the molecular mechanisms governing tumor cell differentiation in MB.

Thyroid hormone (TH) is indispensable for fetal and postnatal brain development, as well as the maintenance of adult brain functions ^9-11^. The major circulating form of TH is thyroxine (T4), which, after de-iodination, transforms into its active counterpart, 3,5,3’-triiodo-L-thyronine, or T3. The actions of TH in the brain are subject to a unique regulatory process involving transport and metabolism. To reach the brain, TH must traverse the blood-brain barrier, facilitated by transporters situated in the membranes of vascular endothelial cells ^12^. Within cells, T3 is primarily responsible for mediating TH’s critical intracellular effects on the transcription of target genes, via interactions between T3 and its nuclear thyroid hormone receptors (TRs) ^13^. In the brain, thyroid receptor α1 (TRα1) predominates and serves as a transcription factor for regulating the transcription of TH target genes, including *Krueppel-like factor 9* (*Klf9*) and *hairless* (*Hr*) ^14^. Patients with MB commonly exhibit hypothyroidism (reduced plasma levels of TH), which is classically considered a treatment-related complication ^15,16^. A possible role of TH, or hypothyroidism, in MB pathogenicity has never been investigated.

During normal development, TH emerges as a critical factor in the differentiation of cells that are the origin of MB, granule neuron precursors (GNPs) ^17^. For example, rats with hyperthyroidism exhibit premature differentiation of cerebellar GNPs, while hypothyroidism impedes GNP differentiation ^18,19^. Pax8 mutant mice, an established model for hypothyroidism, display impaired GNP differentiation, which can be partially rescued through TH treatment ^20,21^. These studies suggest that TH plays an important role in GNP differentiation, prompting us to explore the potential contribution of TH to tumor cell differentiation in MB.

Here, we found that TH plays a pivotal role in MB pathogenicity by promoting terminal differentiation of tumor cells. In the absence of TH (hypothyroidism), the TH receptor, TRα1, is unliganded, allowing it to bind to EZH2 and repress the expression of NeuroD1 by histone methylation of lysine 27 within NeuroD1 regulatory regions. NeuroD1 is a transcription factor that drives terminal differentiation of MB tumor cells ^8,22,23^. Conversely, TH, in the form of T3, counteracts this EZH2-mediated repression of NeuroD1 by disrupting the binding between EZH2 and TRα1, thereby allowing NeuroD1 expression, stimulating terminal differentiation of tumor cells, and reducing MB growth. T3 promotes differentiation and inhibits proliferation of tumor cells from various MB subtypes, including SHH and G3. This suggests that T3-induced differentiation transcends the oncogenic driver mutations present in MB tumor cells. Moreover, T3 exhibits dose-dependent inhibition of MB tumor growth, significantly improves brain tumor symptoms with no observed toxicities in MB mouse models. Together, our results elucidate the mechanisms governing terminal differentiation of MB tumor cells and establish an unprecedented link between TH signaling and MB progression. Our findings offer compelling evidence for T3 as a promising therapeutic modality to promote tumor cell differentiation and suppress MB progression.

## Results

### T3 induces the terminal differentiation of SHH-MB tumor cells

To evaluate a potential role of TH in promoting the terminal differentiation of MB tumor cells, we isolated tumor cells from mice with conditional deletion of *Ptch1* (***Math1-Cre/Ptch1^loxp/loxp^*** mice), an established model for SHH-MB ^24,25^, and treated tumor cells with T3 at different concentrations. Even at the lowest concentration tested (25nM), T3 significantly induced tumor cell differentiation, evidenced by less tumor cell proliferation (EdU labeling), and increased expression of differentiation markers including microtubule-associated protein 2 (MAP2) (**Fig. 1A**) and βIII-tubulin (**Supplementary Fig. 1**). This induction of MB cell differentiation and inhibition of proliferation by T3 was dose-dependent (**Fig. 1B**). No increase in apoptosis (expression of cleaved caspse-3, CC3+) was observed in T3-treated tumor cells (data not shown). These data suggest that T3 treatment suppresses tumor cell proliferation and promotes tumor cell differentiation.

**Figure 1.**
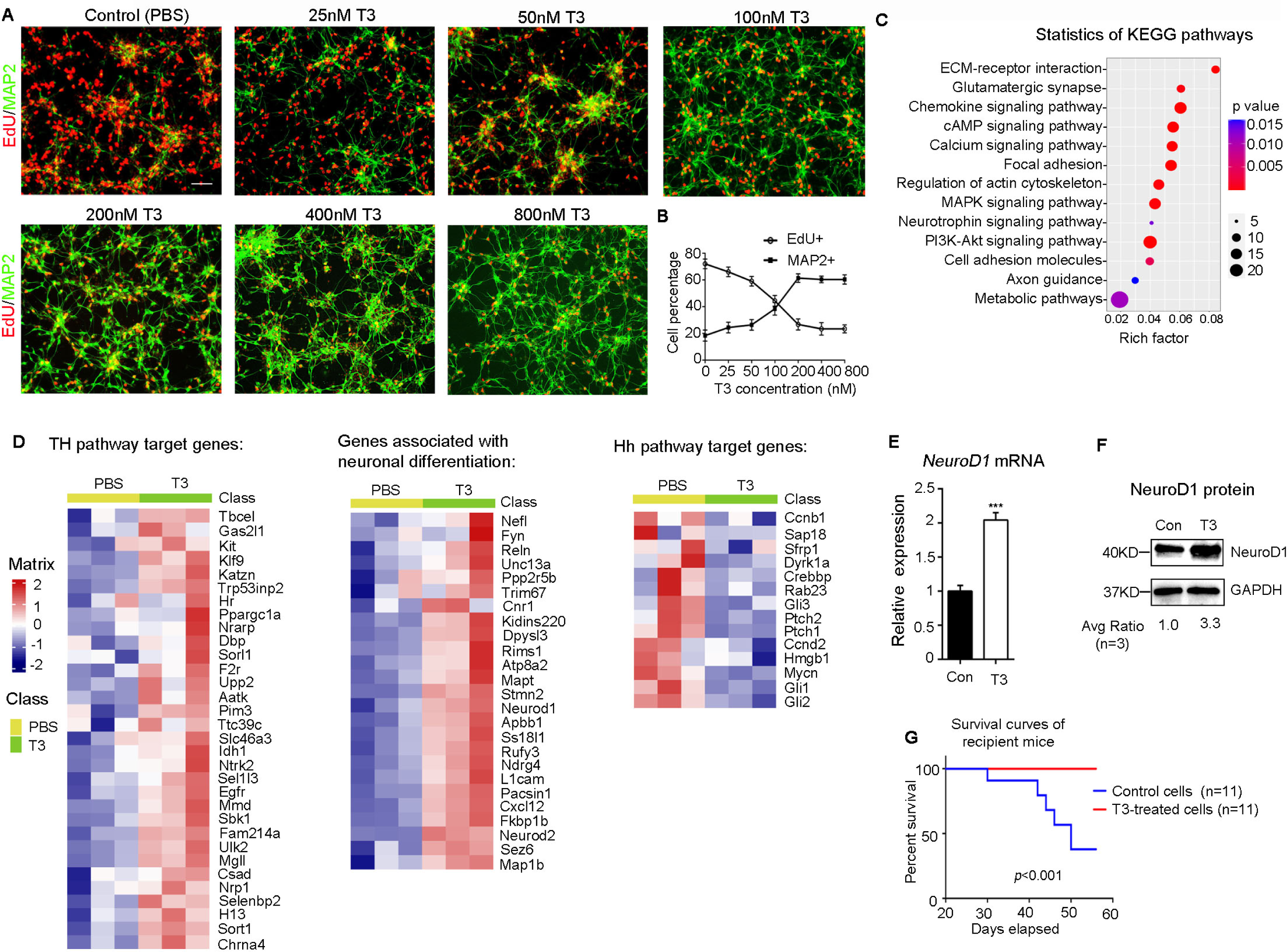
T3 induces terminal differentiation of SHH MB tumor cells. **A-B**, Representative images of MB tumor cells after immunostaining for MAP2 and EdU labelling, following 48 hrs treatment with PBS or T3 (at designated concentrations, **A**). The percentage of MAP2+ cells and EdU+ cells in the culture was quantified, **B**). **C**, KEGG pathways that are upregulated in tumor cells after T3 treatment compared with the control (PBS). **D**, Heat-maps showing expression levels of TH pathway target genes, neuronal differentiation associated genes, and Hh pathway target genes. **E-F**, mRNA and protein expression of NeuroD1 in MB tumor cells treated with PBS (Con) or T3, examined by Q-PCR and western blotting, respectively. **G**, Survival curves of CB17/SCID mice (n=11) after transplantation with T3-treated tumor cells or PBS-treated tumor cells. Scale bar: 50µm (**A**).

To further confirm the terminal differentiation of tumor cells upon T3 treatment, we assessed the transcriptional profiles of tumor cells after treatment with T3 or PBS, by RNA sequencing. We found 4280 transcripts to be differentially expressed between T3- and PBS-treated tumor cells (*p*<0.05, fold changes ≥ 1.5; **Supplementary Table 1**), with 2312 transcripts being upregulated and 1968 downregulated, by T3 treatment compared to PBS controls. KEGG pathway analysis revealed that among the most upregulated pathways in the T3 treated tumor cells were neuronal differentiation-related pathways such as glutamatergic synapse and axon guidance pathways (**Fig. 1C**). Among these upregulated transcripts were TH signaling pathway target genes including *Krueppel-like factor 9* (*Klf9*) and *hairless* (*Hr*) genes ^14^, and neuronal differentiation associated genes such as *Reelin (Reln)* and *Neurofilament light chain (Nefl)* ^26^(**Fig. 1D**), confirming that T3 drives MB tumor cells toward a differentiated phenotype. Among the genes downregulated by T3 treatment were Hh pathway target genes including *Gli1* and *Gli2* (**Fig. 1D**), suggesting that T3 treatment inhibited the pathway that drives tumorigenesis in *Ptch1*-deleted MB cells. NeuroD1, a gene known to regulate the terminal differentiation of MB tumor cells (Cheng et al., 2020b; Zhao et al., 2008) was also upregulated. We confirmed that T3 treatment elevated expression levels of NeuroD1 mRNA and protein in MB cells compared with controls using qPCR and western blot analysis (**Fig. 1E-F**). Importantly, upon intracranial transplantation, T3-treated MB tumor cells failed to develop into tumors in recipient mice (n=11), although untreated tumor cells generated tumors in 6 of 11 recipient mice (**Fig. 1G**). These combined data indicate that T3 induces the terminal differentiation of MB tumor cells and reduces their proliferative capacity and tumorigenic potential.

### Tumor cell differentiation and NeuroD1 transcription are directly repressed by TRα1

To investigate the mechanism underlying T3-induced tumor cell differentiation, we treated tumor cells with 1-850, a specific antagonist of TH receptors that interferes with the interaction between T3 and its receptors ^27^. As shown in **Figure 2A-B**, 1-850 significantly blocked the T3-induced differentiation of tumor cells, indicating that T3 needs to interact with its receptor to induce tumor cell differentiation.

**Figure 2.**
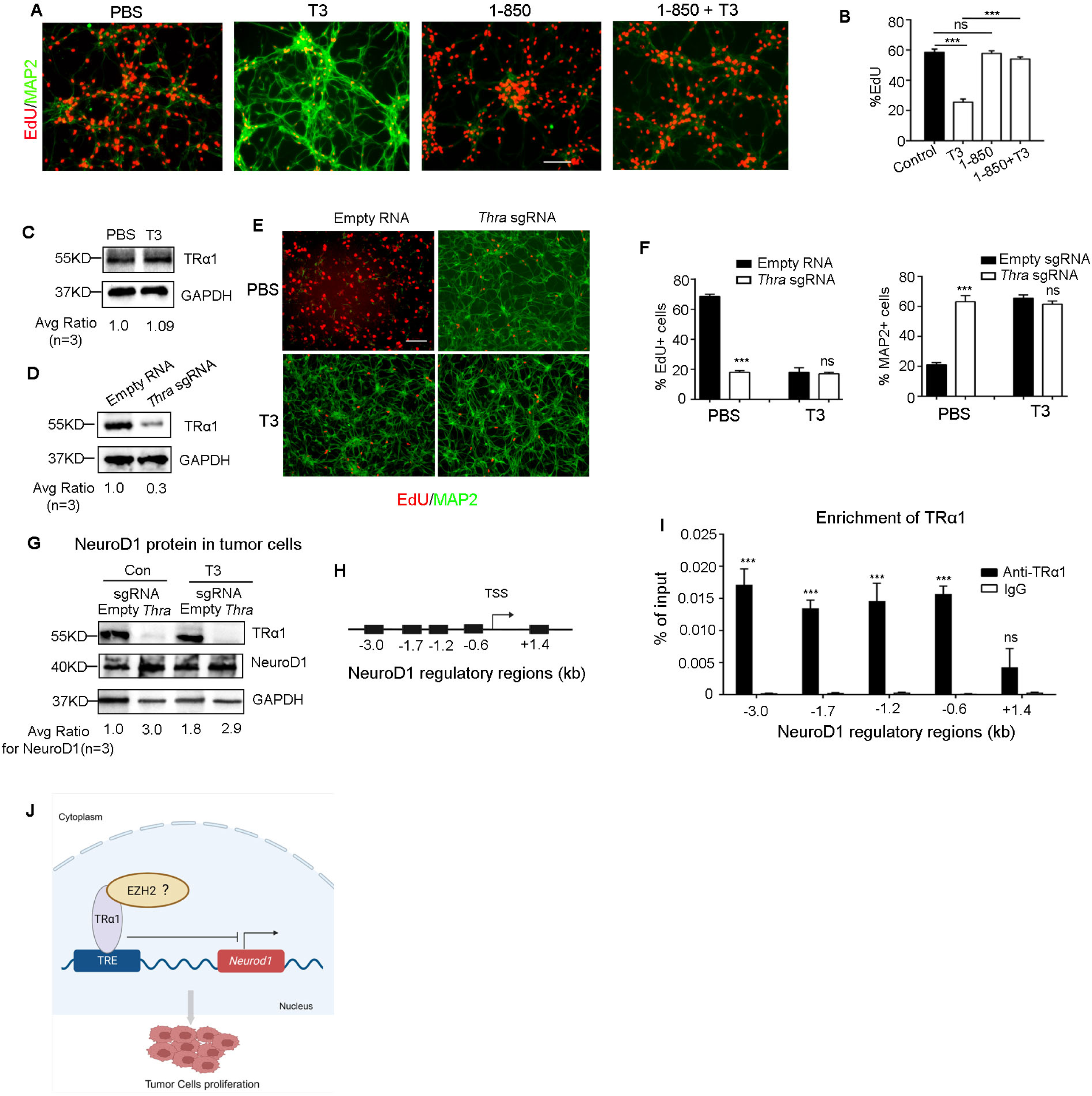
TRα1 mediates T3-induced differentiation of tumor cells. **A-B**, Tumor cells from SHH-MB mice (*Math1-Cre/Ptch1^loxp/loxp^*) were treated with TH receptor-specific antagonist 1-850 or PBS in the presence/absence of 200nM T3, for 48 hrs. After being pulsed with EdU for 2 hrs, tumor cells were harvested to examine differentiation (MAP2+) and proliferation (EdU+) by immunocytochemistry (**A**). Percentages of EdU+ cells in tumor cells (**B**). **C**, Western blot of TRα1 protein in tumor cells from *Math1-Cre/Ptch1^loxp/loxp^* mice, after treatment with T3 (200nM) or PBS for 48 hrs. GAPDH was used as a loading control. **D**, Western blot of TRα1 protein in tumor cells from *CAG-Cas9/Math1-Cre/Ptch1^loxp/loxp^* mice, after being virally infected with an sgRNA specific for *Thra*, or an empty RNA as a control. **E-G**, Immunohistochemistry of tumor cells infected with Thra sgRNA or an empty RNA and treated with T3 or PBS for 48 hrs and pulsed with EdU for 2 hrs, before being harvested to examine proliferation (EdU+) and differentiation (MAP2+) (**E**). Percentages of EdU+ cells or MAP2+ cells (**F**). Western blot using anti-TRα1, anti-NeuroD1, and anti-GAPDH antibodies in tumor cells after *Thra* knockout and treatment with T3 or PBS (**G**). **H**, A schematic showing the distance of established regulatory regions of NeuroD1 gene from the transcription start site (TSS). **I**, ChIP-PCR results showing the enrichment of TRα1 in the NeuroD1 regulatory regions using an antibody against TRα1 or IgG control antibody. Scale bar: 50µm (**A** and **E**). **J**, A schematic showing the repressing role of TRα1 (possibly by binding with EZH2) on the proliferation and NeuroD1 transcription in tumor cells.

The predominant TH receptor in the brain is TRα1, which is a nuclear receptor and functions as a transcription factor that mediates TH signaling by regulating the transcription of target genes ^28,29^. It is encoded by the thyroid hormone receptor alpha (*Thra*) gene. As anticipated, TRα1 is expressed by MB tumor cells and its expression level was not altered by T3 treatment (**Fig. 2C**). To determine whether T3-induced differentiation required the T3 receptor TRα1, we knocked out *Thra* (encoding TRα1) in tumor cells from ***CAG-Cas9/Math1-Cre/Ptch1^loxp/loxp^*** mice. Cells from these mice universally express CRISPR-associated protein 9 (Cas9) ^30^; knockout was achieved by viral transduction with small guide RNAs (sgRNAs) specific for *Thra* or an empty vector as control (**Fig. 2D**). Surprisingly, deletion of *Thra* induced tumor cell differentiation and decreased proliferation, as indicated by a reduction of EdU+ and increase of MAP2+ cells, respectively; the effects of *Thra* deletion were not further impacted by T3 treatment (**Fig. 2E-F**). Similar results were observed after *Thra* knockdown by shRNA (**Supplementary Fig. 2**). These results suggest that MB tumor cell differentiation is intrinsically repressed by TRα1. Knockout or knockdown of *Thra* led to an upregulation of NeuroD1 in tumor cells, which was not further increased by T3 treatment (**Fig. 2G and Supplementary Fig. 2A**). These data suggest that unliganded TRα1 normally inhibits NeuroD1 expression in tumor cells. The binding of TRα1 with the promoter regions of NeuroD1 in tumor cells was confirmed by ChIP-PCR (**Fig. 2H-I**). These results suggest that endogenous TRα1 directly represses NeuroD1 expression in MB tumor cells (**Fig. 2J**).

### T3 interferes with the physical interaction between TRα1 and EZH2

We previously reported that NeuroD1 expression in MB tumor cells is repressed by EZH2-mediated histone 3 lysine 27 trimethylation (H3K27me3) ^8^. Therefore, we examined whether TRα1-suppressed NeuroD1 transcription in tumor cells involved EZH2 and H3K27me3. Comparable levels of EZH2 and H3K27me3 protein were found in tumor cells after T3 or PBS treatment (**Fig. 3A**), suggesting that T3 treatment did not alter the global expression of EZH2 and H3K27me3. However, T3 treatment reduced the enrichment of EZH2 on most of the NeuroD1 promoter regions (except the one located 1.4kb downstream of the transcription start site of NeuroD1) (**Fig. 3B**) in tumor cells compared to the control. These data indicate that T3 treatment reduces the binding of EZH2 to the promoter regions of NeuroD1. Note that T3 treatment did not alter the enrichment of EZH2 on the promoter regions of *Cdkn2a* and *Dio3* in tumor cells (**Fig. 3C**), which are established EZH2-enriched regions ^31,32^. Consistently, T3 repressed the enrichment of H3K27m3 in the promoter regions of NeuroD1 (**Fig. 3D**). These results suggest that T3 regulates NeuroD1 expression by locally interfering with EZH2 enrichment.

**Figure 3.**
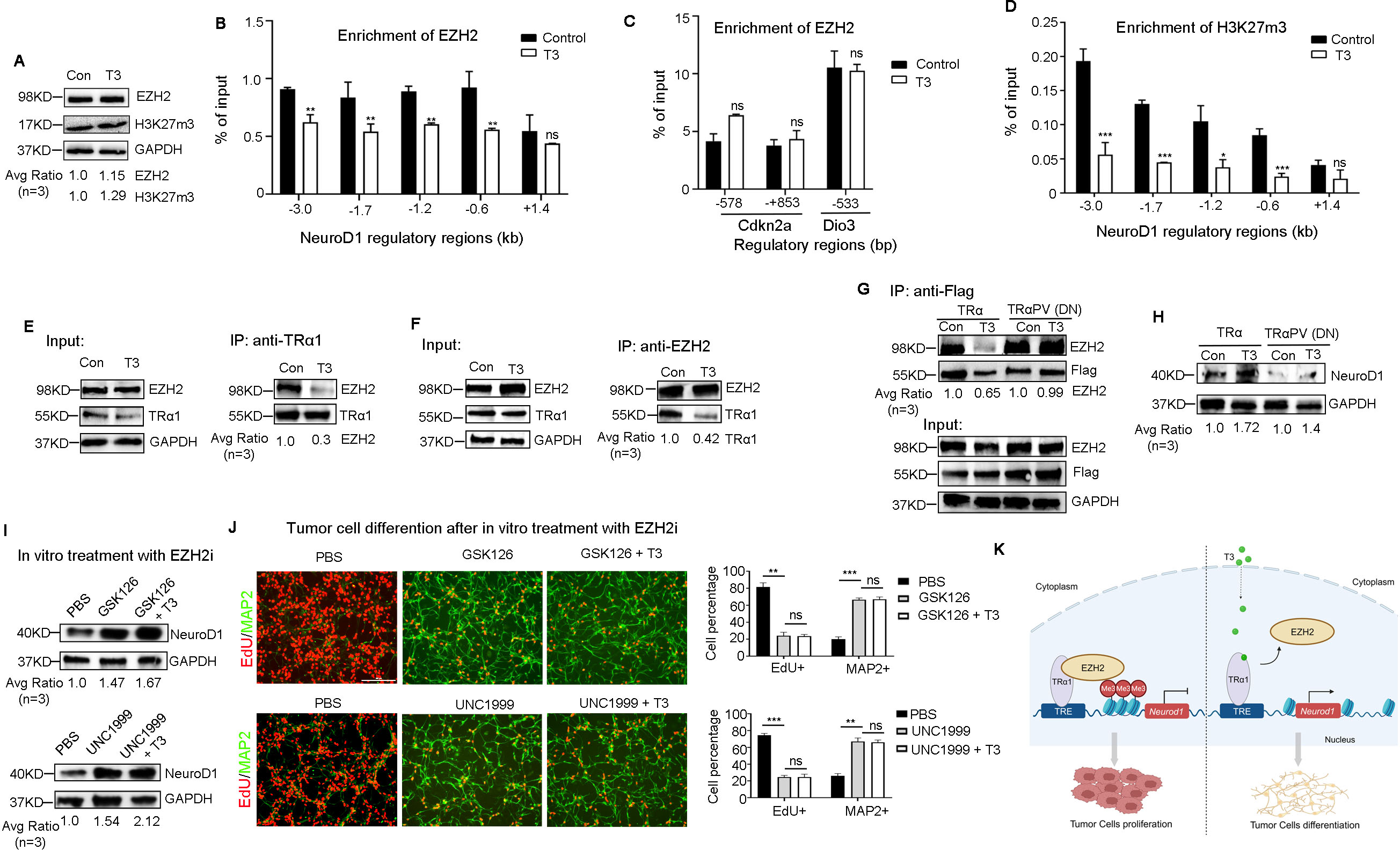
T3 impairs the physical interaction between TRα1 and EZH2, allowing transcription of NeuroD1. **A**, Western blot of EZH2 and H3K27me3 in tumor cells after treatment with T3 or PBS (Con). **B-D**, ChIP-PCR showing effects of T3 or PBS (control) on EZH2 enrichment in NeuroD1 promoter regions (**B**), EZH2 enrichment in promoter regions of established EZH2-enriched genes *Cdkn2a* gene and *Dio3* (**C**), and H3K27me3 enrichment in NeuroD1 promoter regions (**D**) in tumor cells. **E-F**, Co-immunoprecipitation of TRα1 and EZH2 in tumor cells treated with T3 or PBS (Con), using an antibody against TRα1 (**E**) or EZH2 (**F**). **G**, Co-immunoprecipitation of TRα1 and EZH2 in tumor cells transduced with Flag-tagged wild type TRα1 or TRα1PV (dominant negative, lacking T3-interaction domain), using an antibody against Flag. **H**, Western blot of NeuroD1 in tumor cells transduced with wild type TRα1 or TRα1PV. **I-J**, Tumor cells were treated with PBS, GSK126 (1μM) or UNC1999 (1μM) in the absence or presence of T3 (200nM) for 48 hrs before being collected to examine NeuroD1 protein by western blotting (**I**) and tumor cell proliferation and differentiation by immunocytochemistry (**J**). Percentage of proliferative tumor cells (EdU+) and differentiated tumor cells (MAP2+) is shown at the right panel in **J**. **K,** Schematic showing that T3 induces EZH2 expression and promotes terminal differentiation in MB tumor cells by compromising the interaction between TRα1 and EZH2. Scale bar: 50µm (J).

Previous studies had shown that thyroid hormone receptors (such as TRα1) regulate the expression of their target genes by interacting with cofactors such as nuclear corepressor-1^33^ and histone deacetylases ^34^. We then tested for a physical association between EZH2 and TRα1 in tumor cells by co-immunoprecipitation. Indeed, an antibody against TRα1 immunoprecipitated EZH2 and vice versa (**Fig. 3E-F**). Moreover, the association between TRα1 and EZH2 was significantly reduced by T3 treatment (**Fig. 3E-F**). These results suggest that TRα1 and EZH2 physically interact and that such interaction is compromised by T3. To further validate that T3 treatment disturbs the interaction between TRα1 and EZH2, we transduced MB tumor cells with wild type (WT) TRα1 or a dominant negative form of TRα1 (TRα1PV), which lacks the domain for interacting with T3 ^35^. Note that both WT TRα1 and TRα1PV could be immunoprecipitated with EZH2 (**Fig. 3G**), indicating that deletion of the T3 interaction domain from TRα1 did not hinder the ability of TRα1 to interact with EZH2. As anticipated, T3 treatment repressed the interaction of EZH2 with WT TRα1, but not with TRα1PV (**Fig. 3G**), suggesting that T3 needs to bind with TRα1 in order to interfere with the interaction between TRα1 and EZH2. Moreover, in the absence of the TRα1 T3-interaction domain, T3 failed to upregulate NeuroD1 protein expression (**Fig. 3H**), further confirming that T3 induces NeuroD1 expression by binding with TRα1. We next treated tumor cells with EZH2 inhibitors including GSK126 and UNC1999 in the presence or absence of T3, or PBS as a control. As shown in **Fig. 3I-J**, GSK126 and UNC1999 significantly enhanced NeuroD1 expression and induced tumor cell differentiation, phenocopying the T3 treatment. Moreover, T3 treatment failed to further increase NeuroD1 expression or differentiation in GSK126 or UNC1999-treated tumor cells. These data support the notion that T3 induces NeuroD1 expression and differentiation in tumor cells through inhibition of EZH2-mediated H3K27me3. Together, the above findings demonstrate that T3 induces NeuroD1 expression and promotes terminal differentiation in MB tumor cells by compromising the interaction between TRα1 and EZH2 (**Fig. 3K**).

### T3 treatment represses the in vivo growth of SHH-MB with two different driver mutations

Having observed that T3 treatment significantly induces tumor cell differentiation and represses tumor cell proliferation, we next asked whether TH can inhibit the growth of murine SHH-MB in vivo. As SHH-MB can be driven by two different driver mutations (deletion of *Ptch1* or constitutive activation of *Smo*), we tested both. We first treated ***Math1-Cre/Ptch1^loxp/loxp^*** mice with T3 (200ng/g) or PBS as a control (**Fig. 4A**). In control-treated mice, tumor volume progressively increased over 4 weeks. However, in T3-treated mice, no significant increase in tumor volume was observed, indicating that T3 inhibited tumor progression *in vivo* (**Fig. 4B-C**). Consistent with our in vitro data, immunohistochemistry revealed that T3 treatment *in vivo* significantly repressed tumor cell proliferation (Ki67 expression), and enhanced tumor cell differentiation (NeuN expression) (**Fig. 4D**). Based on the expression of cleaved caspase-3 (CC3), no increase in tumor cell apoptosis was detected after treatment with T3 (**Supplementary Fig. 3**). Importantly, T3 treatment significantly prolonged survival of ***Math1-Cre/Ptch1^loxp/loxp^*** mice, compared to control (median survival: T3 treatment, undefined vs PBS treatment, 24 ds, *p*<0.001) (**Fig. 4E**). No obvious reduction in body weight was observed after T3 treatment, compared with PBS treatment (**Fig. 4F**). Consistently, T3 substantially improved the tumor symptoms such as hydrocephalus and ataxia, compared with controls (**Supplementary Videos 1-2**). These data suggest that T3 treatment in vivo induces extensive differentiation of tumor cells and significantly inhibits tumor progression of murine *Ptch1*-deficient SHH-MB.

**Figure 4.**
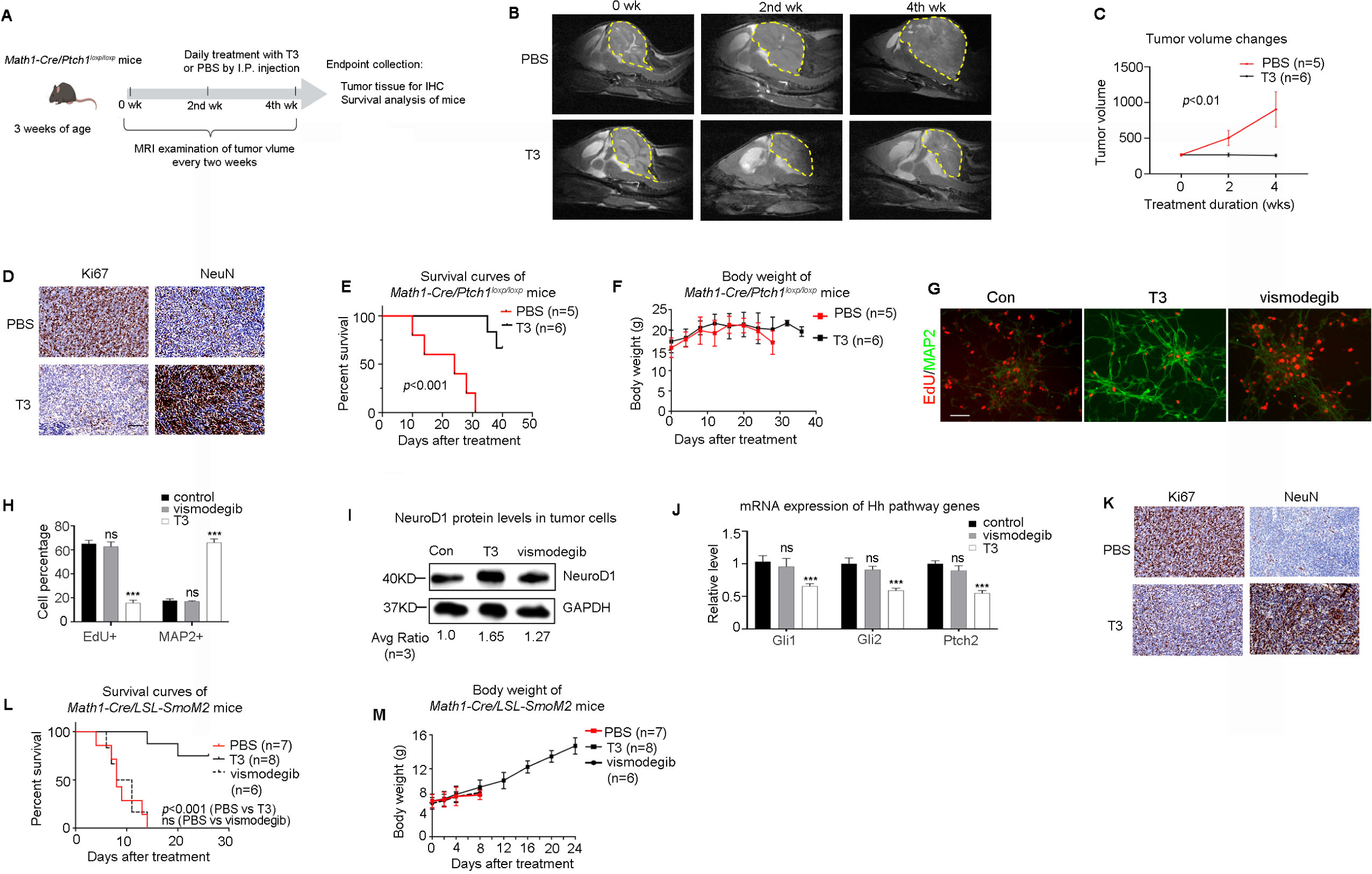
T3 inhibits in vivo growth of murine SHH-MB. **A-C**, A schematic showing the strategy of T3 treatment (**A**). Representative MRI images of brains from the same ***Math1-Cre/Ptch1^loxp/loxp^*** mice before and after treatment with T3 or PBS for 2 weeks and 4 weeks (**B**). Yellow dots indicate tumor area. Tumor volumes are quantified based on MRI images (**C**). **D**, Immunohistochemistry of tumor tissues from ***Math1-Cre/Ptch1^loxp/loxp^*** mice after the T3 or PBS treatment. **E-F**, Survival (**E**) and body weight (**F**) of ***Math-Cre/Ptch1^loxp/loxp^*** mice after treatment with T3 or PBS. **G-H**, Representative immunofluorescence images of tumor cells from ***Math1-Cre/LSL-SmoM2*** mice after treatment with PBS, T3, or vismodegib (**G**). The percentage of EdU+ cells and MAP2+ cells in the culture (**H**). **I-J**, NeuroD1 protein (**I**), and mRNA expression of Hh target genes (**J**) in tumor cells after treatment with PBS, T3, or vismodegib, examined by western blotting and Q-PCR, respectively. **K**, Representative immunohistochemistry images of tumor tissues from ***Math1-Cre/LSL-SmoM2*** mice after treatment with PBS or T3. **L-M**, Survival (**L**) and body weight (**M**) of ***Math-Cre/LSL-SmoM2*** mice after treatment with T3 or PBS. Scale bars: 2mm (**B**); 100µm (**D** and **K**); 50µm (**G**).

We next tested whether T3 produces similar effects in SHH-MB tumor cells derived from constitutive activation of Smo. Smo-driven MB is a ‘drug resistant’ tumor model as this sub-type of MB is resistant to Smo inhibition ^3,36^. These experiments used the established model of constitutive *Smo* activation, ***Math1-Cre/(loxp-stop-loxp)LSL-SmoM2*** mice ^24,25^. First, we asked whether T3 suppressed proliferation and induced differentiation in these cells in vitro. Indeed, T3 significantly suppressed proliferation and stimulated tumor cell differentiation (**Fig. 4G-H**). As expected, constitutive activation of SmoM2 renders tumor cells resistant to killing by vismodegib, an FDA-approved Smo antagonist ^37,38^, and vismodegib failed to cause significant changes in tumor cell proliferation or differentiation (**Fig. 4G-H**). As in *Ptch1*-deficient tumor cells, T3 treatment increased expression of NeuroD1 expression in SmoM2 tumor cells (**Fig. 4I**). Expression levels of *Gli1*, *Gli2* or *Ptch2* mRNAs were significantly reduced by T3 but not by vismodegib, compared with control (**Fig. 4J**), suggesting that T3 treatment inactivated the Hh pathway (both Ptch1 and Smo are in this pathway). In vivo, T3 treatment significantly repressed tumor cell proliferation and increased tumor cell differentiation in ***Math1-Cre/LSL-SmoM2*** mice (as in ***Math1-Cre/Ptch1^loxp/loxp^***mice, **Fig. 4D**) (**Fig. 4K**). No increase in apoptosis was observed in tumor tissues following T3 treatment, compared with PBS (**Supplementary Fig. 3**). T3 significantly prolonged the survival of ***Math1-Cre/LSL-SmoM2*** mice, compared with PBS (median survival: T3 treatment, undefined vs PBS treatment, 8 days; *p*<0.001) (**Fig. 4L**). As expected, vismodegib did not prolong survival (**Fig. 4L**). The body weight of ***Math1-Cre/LSL-SmoM2*** mice steadily increased during T3 treatment (**Fig. 4M**). Even with just 2 weeks of treatment with T3, brain tumor symptoms in ***Math1-Cre/LSL-SmoM2*** mice were markedly alleviated compared with PBS or vismodegib (**Supplementary Videos 3-5**). The above results demonstrate that T3 confers significant tumor inhibitory effects in murine SHH-MB regardless of the driver mutation (*Ptch1* deletion or constitutive Smo activation).

### T3 represses human SHH-MB progression *in vitro* and *in vivo*

We next examined the therapeutic effects of T3 in human SHH-MB tumor cells (ICb-5610MB cells, with inactivating mutations in the *Ptch1* gene) ^39^ and patient-derived orthotopic xenograft (PDOX) models. First, tumor cells were treated with T3, PBS, or cisplatin, a potent DNA damaging agent currently used in MB treatment ^40^ as an additional control. As in the mouse MB cells, T3 treatment of human MB cells significantly decreased the percentage of proliferating cells (EdU+) and significantly increased the number of differentiated cells (MAP2+) (**Fig. 5A, left and center**). More importantly, the number of apoptotic cells after T3 treatment was comparable with that in the PBS-treated control group, indicating that T3 does not compromise the survival of MB tumor cells (**Fig. 5A, right**). As expected, cisplatin also inhibited tumor cell proliferation, but this did not involve the induction of tumor cell differentiation; as also expected, cisplatin caused extensive tumor cell apoptosis (**Fig. 5A**). As was the case in mouse MB tumor cells, in human tumor cells T3 treatment significantly upregulated *NeuroD1* mRNA expression, compared with control or cisplatin treatment (**Fig. 5B**). These combined data indicate that T3 suppresses tumor cell proliferation, induces cell differentiation, and upregulates NeuroD1 in human SHH-MB tumor cells. In addition, in human cells T3 inhibits the Hh pathways as *Gli1* mRNA expression significantly declined after T3 treatment (**Fig. 5C**). Cisplatin reduced expression of both *NeuroD1* and *Gli1* mRNAs, presumably due to the induced apoptosis (**Fig. 5B-C**).

**Figure 5.**
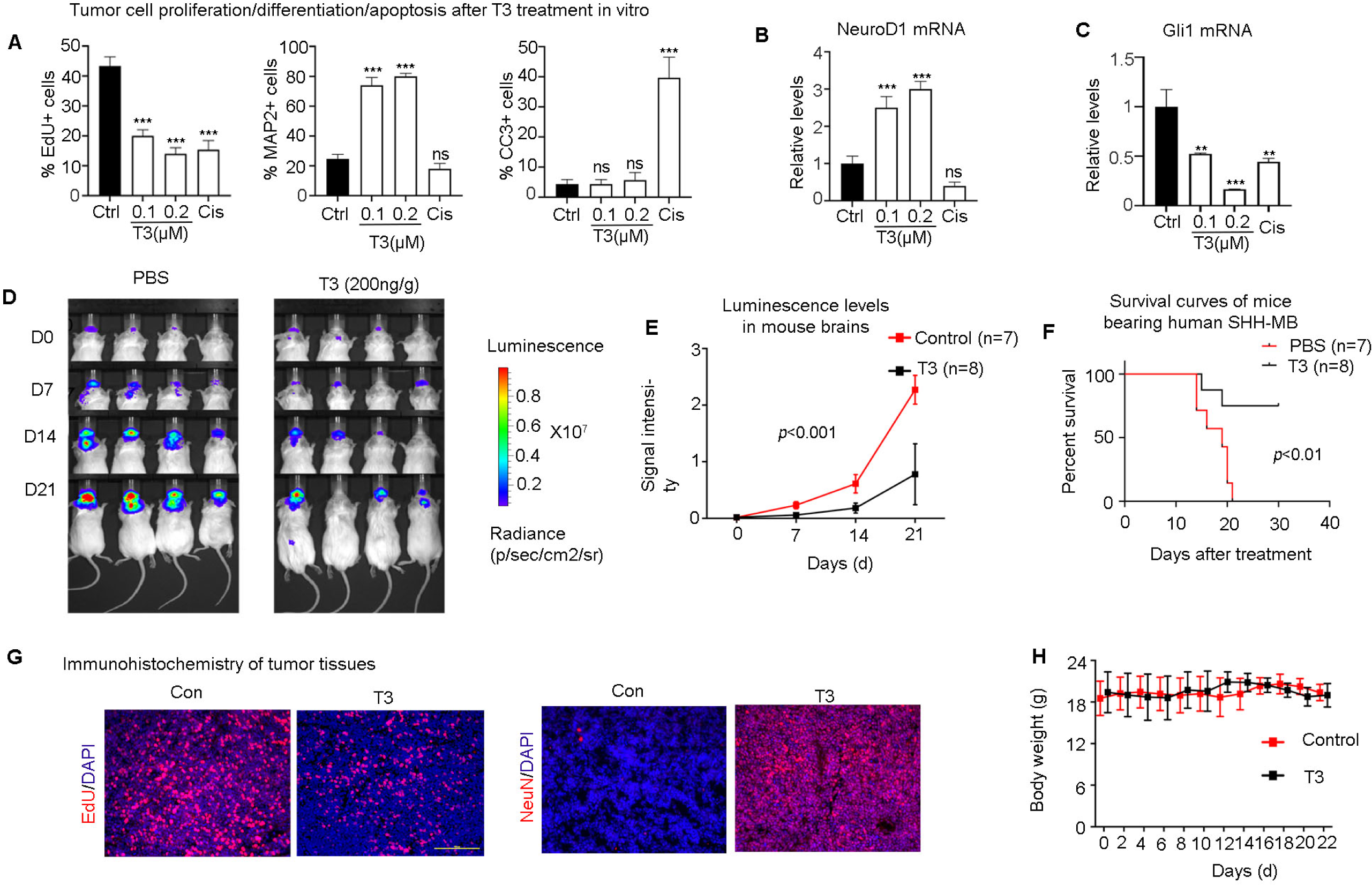
T3 represses the progression of human SHH-MB due to Ptch1 or Smo drivers. **A**, The percentage of proliferative cells (EdU+), differentiated cells (MAP2+) or apoptotic cells (CC3+) in human SHH-MB cells treated with PBS, T3 or cisplatin (Cis) for 48 hrs, based on the immunofluorescence of EdU, MAP2, or CC3. **B-C**, Q-PCR showing the mRNA expression of NeuroD1 (**B**) and Gli1 (**C**) in human SHH-MB cells after treatment with PBS, T3 or cisplatin. **D**, Luminescence images of PDOX-bearing mice bearing SHH-MB cells, after treatment with PBS or T3 from day 0 (D0) to day 21 (D21). **E,** Quantification of brain luminescence intensity. **F,** Survival curve of PDOX-bearing mice after treatment with PBS or T3. **G,** Representative immunofluorescence images of frozen tissue sections from PDOX-bearing mice after control or T3 treatment; EdU was administered 2 hrs before sacrifice. Right panels show staining for NeuN (human differentiation marker) DAPI was used to counterstain cell nuclei. **H**, body weight of PDOX-bearing mice during treatment with T3 or PBS. Scale bar: 100µm (**G**).

We then assessed the effects of T3 on tumor growth in patient-derived orthotopic xenograft (PDOX) mouse models. Human ICb-5610MB cells transduced with a luciferase expressing construct were transplanted into the cerebella of *CB17/SCID* mice. After tumors were established, mice were daily treated with PBS (control) or T3 (200ng/g) by I.P. injection. After 3 weeks of treatment, tumor progression in the T3-treated mice was significantly inhibited compared with controls (**Fig. 5D**). Quantification of luciferase levels in the brain showed that tumors progressively increased in control-treated mice, but the rate of increase was significantly reduced by T3 treatment (**Fig. 5E**). T3 treatment also significantly improved survival (median survival: T3 treatment, undefined vs PBS treatment, 19 days; *p*<0.01. **Fig. 5F**). Consistent with our observations in murine tumor tissue, T3 treatment inhibited tumor cell proliferation and promoted tumor cell differentiation in PDOXs of SHH-MB (**Fig. 5G**). No increase in apoptosis was observed following T3 treatment, compared with PBS (**Supplementary Fig. 3**). Moreover, body weight was comparable following T3 or control treatment (**Fig. 5H**), suggesting that T3 treatment did not produce obvious toxicities in PDOX SHH-MB mice. These data demonstrate that T3 potently induces tumor cell differentiation and inhibits the growth of human SHH-MB in mice.

### T3 inhibits the progression of Group 3 MB

Group 3 (G3) is the most aggressive form of MB. We therefore examined whether T3 induces differentiation of tumor cells from G3-MB, asked if increased expression of NeuroD1 by T3 is a key mechanism, and tested the efficacy of T3 treatment in G3-MB Patient-Derived Orthotopic Xenografts (PDOXs). As there is no genetic mouse model of G3 MB, our *in vitro* and *in vivo* assays used human MB tumor cells. First, we treated RCMB20 or RCMB28 cells, two established PDOX lines of human G3-MB ^41^, with T3 or PBS (**Fig. 6A**). T3 significantly reduced the proliferation of both RCMB28 and RCMB20 cells and significantly enhanced differentiation, compared with control (**Fig. 6B-C**). As in SHH-MB tumor cells, no increase in apoptosis was observed in G3-MB tumor cells after T3 treatment, compared with control (data not shown). The inhibitory effect of T3 on the proliferation of RCMB28 cells was confirmed by tumoroid forming assays (**Supplementary Fig. 5**). T3 also elevated the amount of NeuroD1 protein in tumor cells indicating that T3 induces the expression of this essential neuronal differentiation protein in G3-MB tumor cells (**Fig. 6D**).

**Figure 6.**
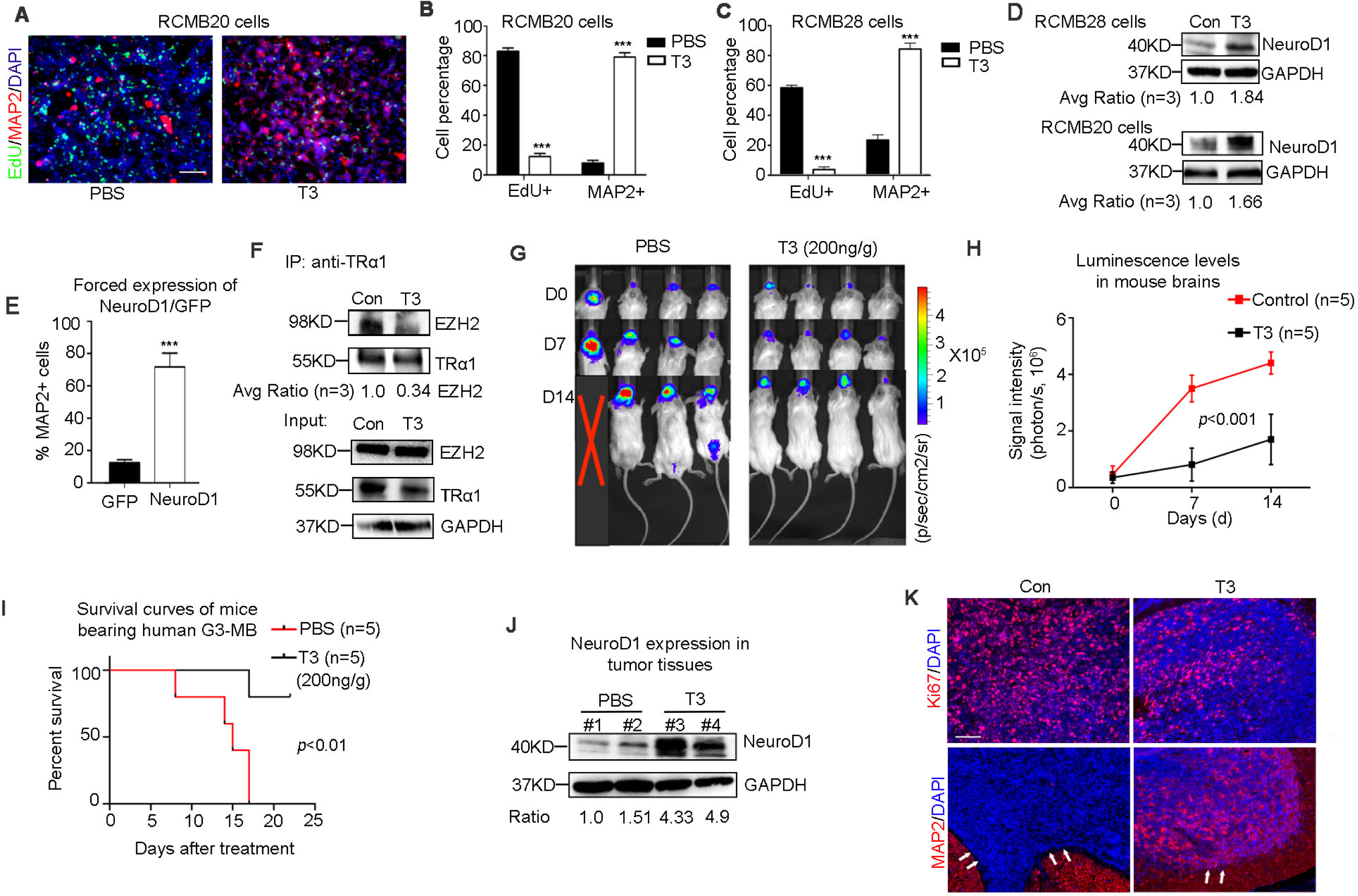
T3 inhibits the pathogenicity of human G3-MB. **A-C**, Representative immunofluorescence images of RCMB20 cells after treatment with T3 or PBS after immunostaining for EdU and MAP2 (**A**). DAPI was used to counterstain cell nuclei. Percentages of EdU+ cells and MAP2+ cells in the culture of RCMB20 (**B**) and RCMB28 cells (**C**). **D**, Western blot showing increased NeuroD1 expression in RCMB28 cells and RCMB20 cells, after T3 treatment for 48 hrs. **E**, Percentage of MAP2+ cells in RCMB28 cells after transduction of GFP or NeuroD1-GFP. **F**, Co-immunoprecipitation of EZH2 and TRα1 in RCMB28 cells after PBS or T3 treatment, using an antibody against TRα1. **G-I**, Luminescence images of PDOX mice bearing RCMB28 cells (**G**), brain luminescence intensity (**H**) and survival (**I**) of PDX mice after PBS or T3 treatment. **J**, Western blot showing increased NeuroD1 expression in tumor tissues from 2 of the animals shown in panel G (#1-#4), after PBS or T3 treatment. **K**, Representative immunohistochemistry image of Ki67 and MAP2 in frozen sections (**L**) from G3-MB bearing mice after PBS or T3 treatment. Note that arrows indicate tumor tissues in **K**. DAPI was used to counterstain cell nuclei. Scale bars: 50µm (**A**); 100µm (**K**).

Second, we tested if this increased expression of NeuroD1 by T3 could be a mechanism responsible underlying the T3-induced decrease in proliferation and increase in differentiation in MB. We transduced RCMB28 cells with an expression vector carrying either NeuroD1-GFP or GFP alone as a control. Forced expression of NeuroD1 significantly promoted tumor cell differentiation, as evidenced by increased numbers of MAP2+ cells in NeuroD1-transduced cells compared with control cells (**Fig. 6E**), suggesting that the enhanced expression of NeuroD1 by T3 does drive differentiation of G3-MB tumor cells. In G3-MB cells, as in SHH-MB cells (both driver mutation), intrinsic TRα1 interacts with EZH2, and this interaction was impaired by T3 treatment (**Fig. 6F**). These data suggest that T3 induced NeuroD1 expression in G3-MB tumor cells, by a similar mechanism as in SHH-MB tumor cells.

Finally, we generated G3-MB PDOXs by intracranial transplantation of RCMB28 cells transduced with a luciferase expressing construct. After tumors were established (∼ 21 days), mice were treated with T3 or PBS by I.P. injection. T3 treatment significantly slowed tumor progression (**Fig. 6G**) and reduced the luciferase levels in mouse brains compared with PBS (**Fig. 6H**). Importantly, T3 significantly prolonged survival (median survival: T3 treatment, undefined vs PBS treatment, 15 days; *p*<0.01. **Fig. 6I**). As in the human SHH-GB PDOX model, T3 increased the amount of NeuroD1 protein in PDOX tissues (**Fig. 6J**), suggesting that T3 treatment induces NeuroD1 expression in G3-MB tumor cells. As in SHH-MB, T3 repressed proliferation and enhanced differentiation of tumor cells in tumor tissue (**Fig. 6K**). These results demonstrate that T3 treatment induced the differentiation of human G3-MB tumor cells and inhibited G3-MB growth in a PDOX model. Importantly, our finding that T3 promoted tumor cell differentiation and suppressed tumor growth in both SHH-MB and G3-MB indicates that the tumor inhibitory effects of T3 are not limited to particular oncogenic mutations in MB tumor cells.

### T3 treatment causes no toxicities in mice

Having observed the promising effects of T3 in inhibiting MB growth, we examined possible toxicities of T3 in mice. The recommended dosage of T3 for patients is 75-100µg/day ^42^. Converting this to mice (based on body surface area) ^43^, the equivalent dose is 10-20ng/g/day. We therefore treated ***Math1-Cre/Ptch1^loxp/loxp^*** mice with 4 different doses of T3 (2ng/g-body weight, 20ng/g, 200ng/g and 2µg/g) or PBS, by I.P. injection daily (**Fig. 7A, n=3**) for 3 weeks. No mice died during the 3-week treatment. T3 levels in brain increased in a dose-dependent manner, confirming that I.P. injection represents an effective approach to deliver T3 to the brain (**Fig. 7B**). Note that by the end of 3-week T3 treatment, plasma levels of free T3 did not differ significantly between T3 and PBS treated mice (except 2µg/g) (**Fig. 7C**), possibly due to the intricate TH homeostatic mechanisms that have evolved to regulate systemic T3 levels ^44,45^.

**Figure 7.**
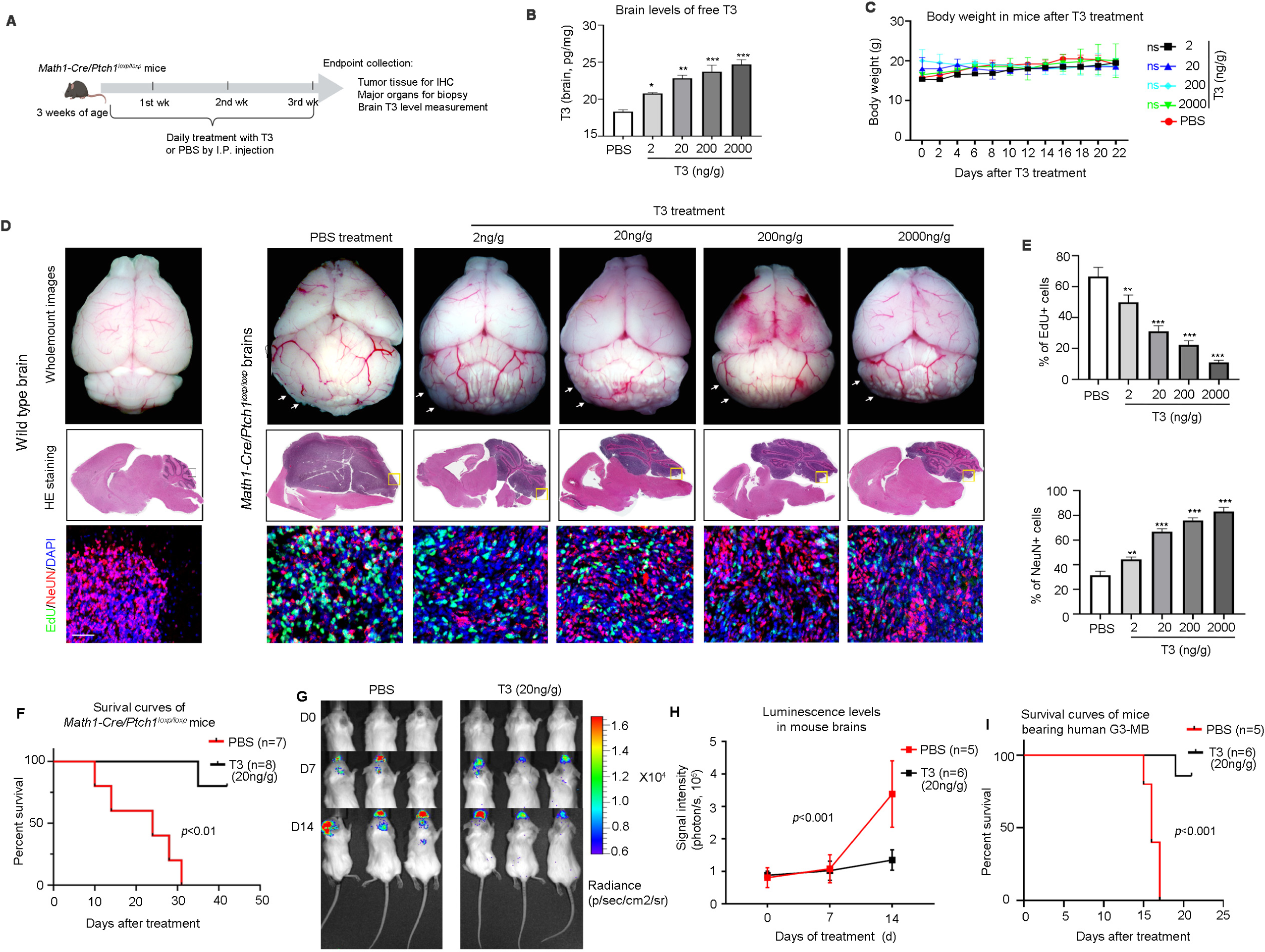
T3 does not cause toxicity in MB-bearing mice. **A**, Schematic for T3/PBS toxicity study in ***Math1-Cre/Ptch1^loxp/loxp^*** mice. **B-C**, T3 levels in the tumor tissue (**B**) and plasma (**C**) in mice after T3/PBS treatment for 3 weeks. **D-F**, Measures of food consumption (**D**), water consumption (**E**) and body weight (**F**) of mice during the T3/PBS treatment. **G-H**, Heart rate measured weekly (**G**), and coefficient of variation (CV) of heart rate (**H**) of mice throughout the 3 weeks of T3/PBS treatment, based on the ECG recordings. **I**, Representative images of whole-mount brains (upper panel), and HE staining (middle panels) and corresponding (boxed regions) immunohistochemistry (lower panels) of brain sections, from wild type or ***Math1-Cre/Ptch1^loxp/loxp^*** mice after treatment with PBS or indicated doses of T3. Arrows pointing to tumors in mouse cerebella. **J,** Quantification of percentages of proliferating cells (EdU+) or differentiated cells (NeuN+) in tissues sections as in panel I. **K**, Survival curves of ***Math1-Cre/Ptch1^loxp/loxp^*** mice after treatment with 20ng/g T3 or PBS, once a day, by I.P injection. **L,** Luminescence images of recipient mice transplanted with RCMB28 cells, after the treatment with 20ng/g T3 or PBS by I.P. injection, once a day. **M,** Quantification of brain luminescence intensity. **N,** Survival curves of PDOX mice after PBS or T3 treatment. Scale bars: 100µm (**I**).

The classic signs of thyrotoxicosis in mice are increased food/water intake, weight loss, and tachycardia. We therefore evaluated the following potential effects of T3 in mice (n=3) over a 3-week treatment period: 1) food/water consumption; 2) body weight; 3) cardiac electrophysiology; 4) hematologic analysis; and 5) histopathology of major organs. Food and water consumption in T3-treatment mice was elevated above control levels during the 1^st^ and 2^nd^ week of T3 treatment. However, food/water consumption declined to levels close to that in controls as the treatment progressed (**Fig. 7D-E**). No significant changes in body weight were observed in mice after T3 or PBS treatment (**Fig. 7F**), suggesting that the treatment with T3 (2ng/g-2µg/g) did not cause weight loss. We next assessed cardiac electrophysiology in mice after T3 or PBS treatment. Electrocardiogram (ECG) recording showed that all mice had normal sinus rhythm, with no arrhythmias detected during T3 treatment (data not shown). Heart rate (HR) in PBS-treated mice declined during the treatment (**Fig. 7G**), likely due to the intracranial hypertension caused by tumor progression ^46^. HR in T3-treated mice increased (∼15%) at all doses (**Fig. 7G**). After completion of the T3 treatment, no significant difference in HR was observed in the T3-treated mice (**Fig. 7G**). We also determined the coefficient of variance (CV) of HR over the 3 week treatment to assess HR variability over time ^47^. No significant difference in the CV was detected among T3-treated mice, or between T3-treated and PBS-treated mice (**Fig. 7H**), together suggesting that T3 treatment did not significantly affect HR in mice. These combined data demonstrate that T3 treatment ranging from 2ng/g-2µg/g causes no significant toxicities in mice.

No abnormalities in the hematology parameters were observed in mice after T3 treatment (**Supplementary Table 2**). No alterations were detected in the biopsy and histological analyses of major organs including thyroid gland, lung, liver, heart, spleen, kidney and gastrointestinal track (data not shown). These data suggest that T3 ranging from 2ng-2µg/g-body weight causes no toxicities in MB-bearing mice.

### T3 at a clinically-relevant dose increases survival in Ptch1 and G3-MB mice

We first examined the effects of the increasing doses of T3 on tumor growth in ***Math1-Cre/Ptch1^loxp/loxp^***mice by histological analyses and immunohistochemistry. Consistent with our previous report ^24^, in PBS-treated mice cerebellar structure was substantially disrupted due to the tumor growth, compared with wildtype mice of the same age (**Fig. 7I**). However, in T3 treated mice, tumor size was reduced in a dose-dependent manner by all T3 doses including 2ng/g, compared with the control (PBS) (**Fig. 7I**). Moreover, cerebellar structure was partially restored by T3 treatment, especially at the higher doses (**Fig. 7I**). The tumor symptoms significantly improved with T3 treatment, even at the lowest dose of 2ng/g (**Supplemental Video 6-10**). We then examined tumor cell differentiation and proliferation markers by immunohistochemistry in tumor sections. Consistent with the gross pathology findings, higher doses of T3 were associated with increasing numbers of NeuN+ (human neuronal differentiation marker) cells and declining numbers of EdU+ cells (**Fig. 7I-J**), suggesting that T3 treatment induced tumor cell differentiation and repressed tumor cell proliferation in a dose-dependent manner.

Finally, we tested the tumor inhibitory effects of T3 at 20ng/g, a dose equivalent to the typical human dose (75-100µg per day). ***Math1-Cre/Ptch1^loxp/loxp^***mice were treated with 20ng/g T3 or PBS by I.P. injection, once a day. The T3 treatment significantly prolonged the survival of ***Math1-Cre/Ptch1^loxp/loxp^***mice compared to the control treatment (median survival: T3 treatment, undefined vs PBS treatment, 24 days; *p*<0.01. **Fig. 7K)**. In the aggressive G3-MB, 20ng/g T3 treatment significantly inhibited the in vivo progression of human G3-MB tumor cells (**Fig. 7L-M**) and improved the survival of the PDOX-bearing mice (median survival: T3 treatment, undefined vs PBS treatment, 16 days; *p*<0.001. **Fig. 7N**). These data confirm the therapeutic efficacy of T3 at a clinically relevant dose, in MB treatment.

The above combined data indicate the T3, even at doses 100x typical human dosing, do not produce obvious toxicities in mice, and confirm the efficacy of T3 in repressing tumor progression, inducing tumor cell differentiation, and improving MB symptoms. Our findings demonstrate that T3 represents a safe and effective approach to treat MB.

## Discussion

Despite the established use of conventional chemotherapy and radiation as primary treatments for many malignancies, these approaches often come with challenges such as drug resistance and systemic toxicity, which limit their therapeutic efficacy. An alternative strategy known as “Differentiation therapy”, offers a unique and potentially less toxic approach to cancer treatment. This approach aims to induce terminal differentiation in cancer cells, leading to a loss in their proliferative and tumorigenic capacity ^48,49^. However, the mechanisms governing terminal differentiation of cancer cells remain poorly understood across various malignancies, hindering the development of differentiation-based treatment strategies.

In this study, we shed light on the potential of TH to induce terminal differentiation in MB tumor cells by promoting NeuroD1 transcription. Our findings reveal that TH-driven tumor cell differentiation leads to a substantial reduction in tumor cell proliferation and effectively inhibits MB growth in vivo. Importantly, the striking tumor-inhibitory effects of TH extends to both SHH-MB and G3-MB, suggesting that T3 therapy could have broad application across MB subgroups. This study establishes T3 treatment as an innovative and promising strategy for MB therapy, highlighting the advantage of differentiation-based strategies in cancer treatment. T3, an FDA-approved drug for hypothyroidism, has demonstrated ability to cross the blood-brain barrier ^50,51^. Our studies also establish that T3, at clinically relevant doses, can penetrate the brain and suppress tumor growth without any apparent toxicities (even at 2µg/g). Given the favorable safety profile of T3, our findings provide a compelling rationale for investigating T3 in clinical trials for MB treatment, either as a stand-alone therapy or in combination with other treatment modalities.

Hypothyroidism is a prevalent endocrine disorder in patients with MB, primarily considered a consequence of aggressive tumor treatments, particularly craniospinal irradiation ^15,16^. However, the potential impact of hypothyroidism on MB progression has remained unexplored. Our studies present the first compelling evidence that TH signaling actively inhibits MB progression, suggesting that hypothyroidism might actually contribute to MB pathogenesis. Our findings underscore the importance of routinely assessing TH levels and hypothyroidism in MB patients. Furthermore, the widespread occurrence of hypothyroidism among MB patients offers additional support for the idea of using T3 as part of MB treatment.

It has been known for decades that TH signaling plays a pivotal role in regulating normal cerebellar development. Both in rodents and humans, hypothyroidism has been linked to abnormal morphogenesis and functional alterations in the cerebellum. Normal differentiation of cerebellar granule neuron precursors (GNPs) is disrupted in mice by altered TH levels ^18,19^, deficiency in TH receptors ^52^ or mutations in deiodinase ^53^. The above evidence emphasizes the significance of TH signaling in promoting the differentiation of neuronal progenitors, which prompted us to investigate the potential functions of TH in MB tumor cell differentiation.

Despite these longstanding insights, the precise molecular mechanisms underlying the critical role of TH in cerebellar development remain incompletely understood. Notably, only a limited number of T3 target genes in neuronal cells have been described thus far. Our studies demonstrate that T3 stimulates tumor cell differentiation through directly inducing NeuroD1 transcription. Our ChIP-PCR experiments further confirm that NeuroD1 serves as a target gene of T3 in MB tumor cells. Given the established role of NeuroD1 in the differentiation of cerebellar GNPs and MB tumor cells, our studies identify NeuroD1 as a novel T3 target gene during MB tumor formation, and likely in normal cerebellar development.

In our mechanistic investigations, we found that knockout or knockdown of TRα1 significantly upregulated NeuroD1 expression and promoted differentiation in tumor cells. These findings suggest that TRα1 acts as a repressor of NeuroD1 expression and terminal differentiation in MB tumor cells. To accomplish this, TRα1 recruits EZH2 as a corepressor to inhibit NeuroD1 transcription by mediating the histone methylation at lysine 27 (K3K27m3) within NeuroD1 promotor regions. This mechanism is further supported by our observations that pharmaceutical inhibition of EZH2 mirrored the effects of TRα1 knockout in stimulating NeuroD1 expression and inducing differentiation in tumor cells. Previous studies had shown that EZH2 engages in gene-specific chromatin modification by interacting with transcription factors such as estrogen receptor and NF-kB ^54,55^. In our studies, T3 treatment disrupts the physical interaction between EZH2 and TRα1 in tumor cells, thereby inducing NeuroD1 expression by counteracting histone methylation. It is noteworthy that global levels of EZH2 and H3K27me3 expression remained unaffected in tumor cells following T3 treatment, suggesting that T3 regulates NeuroD1 expression in a gene-specific manner. One intriguing question arising from our research concerns how T3 treatment alters the binding of TRα1 with EZH2. Prior studies based on the crystal structures of TRα1, have shown that T3 induces ligand-dependent conformational changes in TRα1, affecting its ability to interact with co-repressors or co-activators ^56,57^. Indeed, in the presence of T3, TRα1 loses its capacity to form homodimers or heterodimers with RXR ^58^. Therefore, it is plausible that T3 treatment may result in conformational changes of TRα1, impairing its interaction with EZH2 in tumor cells. Future studies are warranted to confirm this. Collectively, these findings establish the TH/TRα1/EZH2/NeuroD1 signaling axis and its pivotal role in promoting tumor cell differentiation and MB pathogenesis.

## Materials and Methods

### Mice

*Math1-Cre* mice, *Ptch1^loxp/loxp^* mice, *LSL(<u>L</u>oxp-<u>S</u>TOP-<u>L</u>oxp)-SmoM2* mice, *Math1-CreER* mice, *CAG-Cas9* mice were purchased from the Jackson Laboratory. All mice were bred and genotyped as recommended by the Jackson Laboratory. *CB17/SCID* mice were bred in the Fox Chase Cancer Center Laboratory Animal Facility (LAF). All animals were maintained in the LAF at Fox Chase Cancer Center and all experiments were performed by procedures approved by the Fox Chase Cancer Center Animal Care and Use Committee.

### Plasmids and chemicals

T3 (#16028) or 1-850 (#17248) was purchased from Cayman Chemical Company. Vismodegib (#S1082) was purchased from Selleck Chemicals LLC. Vectors including pCMV-shRNA-TRα1 and pU6-sgRNA#1TRα1-sgRNA#2TRα1, were generated by Vectorbuilder. Plasmids containing pMSCVhygro-Ezh2 (#24926) or pMSCVhygro-Ezh2-F667I (#24927) were purchased from Addgene. Vectors encoding wild type TRα1 (pCMV-Flag-TRα1) or mutant TRα1 (pCMV-Flag-TRα1PV) were kindly provided by Dr. Sheue-yann Cheng Laboratory, National Cancer Institute, Bethesda, MD.

### Tumor cell culture and treatment

Primary cells were freshly isolated from tumor-bearing mice at 6 to 8 weeks of age, as previously described^59^. Briefly, tumor tissues were isolated from mouse brains and digested in a papain solution to obtain a single cell suspension and then centrifuged through a 35% and 65% Percoll gradient. Cells from the 35% to 65% interface were suspended in Dulbecco’s PBS (DPBS) plus 0.5% BSA. Cells were then suspended in NB-B22 (Neurobasal medium with B22 supplement without T3, and the B22 supplement was prepared following the protocol generated in Jacob Hanna’s lab^60^). Cell suspension was plated on poly-D-lysine (PDL)-coated coverslips (BD Biosciences). In most in vitro experiments, 200nM T3 (unless other concentrations designated) was added to the culture medium, and 200nM vismodegib or 1µM cisplatin was used as controls. After treatment for 48 hrs, tumor cells were harvested to examine their proliferation, differentiation or apoptosis by immunocytochemistry. DAPI was used to counterstain cell nuclei for quantifying cell percentage.

For tumoroid culture, human MB cells were plated in a 6-well plate (2×10^5^ cells/well) using NB-B22 medium supplemented with 10ng/ml FGF and 20ng/ml EGF for 4-5 days, in the presence/absence of 200nM T3. For counting the number of tumoroids, 50μl of tumoroid suspension were collected and the number of tumoroids was counted under a microscope. Established tumoroids were defined by a minimal sphere size spanning at least 50μm in diameter.

### Lentivirus preparation

293FT cells or Lenti-X 293T cells (carrying pLVX lenti-backbone) were transfected with the plasmid using the calcium phosphate transfection kit (Thermo Fisher). Following the transfection, culture medium was replaced with DMEM plus 10% FBS and 2-5 mM Sodium Butyrate to increase virus production efficacy. The cells were then cultured for 36 hrs before culture medium was harvested. The culture medium was then concentrated by ultracentrifugation and stored at -80°C.

### Gene knockdown or knockout

To knock down *THRA* in tumor cells, lentivirus carrying shRNA constructs targeting *THRA* or scrambled shRNA (Vectorbuilder) was generated. Tumor cells were virally transduced with the shRNA vectors, and validated for *THRA* knockdown in tumor cells by examining TRα1 through western blotting. All sequences for shRNAs were listed in **Supplementary Table 3**.

To knock out *THRA* in tumor cells, we crossed *CAG-Cas9* mice with *Math1-Cre/Ptch1^loxp/loxp^* mice to generate *CAG-Cas9/Math1-Cre/Ptch1^loxp/loxp^* mice in which all tumor cells express Cas9. A lentiviral vector encoding mcherry tagged small guide RNAs (sgRNAs) targeting *THRA*, was purchased and used for virus preparation. Note that this lentiviral vector contains two guide RNAs targeting *THRA*, to enhance the knockout efficiency. Tumor cells were virally transduced with sgRNAs specific for *THRA*, or an empty vector as a control. At 48 hrs following the infection, infected cells were purified by FACS based on mcherry expression. Western blotting was performed to examine levels of TRα1 protein in infected cells, to validate the deletion of *THRA*. Sequences of sgRNAs for *THRA* knockout were included in **Supplementary Table 3**.

### Immunostaining, western blotting and co-immunoprecipitation (Co-IP)

Immunofluorescence staining of sections and cells was carried out according to standard methods. Briefly, sections or cells were blocked and permeabilized for 1 hr with PBS containing 0.1% Triton X-100 and 10% normal goat serum, stained with primary antibodies overnight at 4°C and incubated with secondary antibodies for overnight. Sections or cells were counterstained with DAPI and mounted with Fluoromount-G (Southern Biotech) before being visualized using a Nikon Eclipse Ti microscope. In some experiments, tumor cells were pulsed with EdU (0.02µg/µl) for 2 hrs prior to immunostaining. Primary antibodies used for immunofluroscence assay include MAP2 (1:500, Millipore, Ab5622); NeuN (1:200, Abcam, Ab104224); Cleaved caspase-3 (1:500, CST, 9664S).

For western blot analysis, cells were lysed in RIPA buffer (Thermo) supplemented with protease and phosphatase inhibitors (Thermo). Total lysate containing equal amount of protein were separated by SDS-PAGE gel and subsequently transferred onto PVDF membrane. Membranes were then subjected to probe with antibodies. Western blot signals were detected by using SuperSignal West Pico Chemiluminescent substrate and visualized using a iBright imaging system (Model CL1500, ThermoFisher). Primary antibodies used in this study include NeuroD1 (1:1000, Abcam, Ab60704); TRα1 (1:1000, Abcam, Ab53729); EZH2 (1:1000, CST, 5246); GAPDH (1:1000, CST, 2118S). Protein levels were quantified from 3 experiments using ImageJ and normalized to GAPDH.

Co-IP experiments were performed using the DynabeadsTM Protein G Immunoprecipitation Kit according to the manufacturer’s instruction. Briefly, cells were homogenized and incubated on ice for 15 mins, which were then centrifuged to remove the cell debris. 50 µl cell lysate was used as an input, which was resuspended with Dynabeads™ magnetic beads. After incubation for 9-12 hrs at 4°C, the beads were washed thoroughly with provided washing buffer. The beads were then eluted using the elution buffer, which was heated for 10 mins at 70°C before being used for western blotting.

### Quantitative polymerase chain reaction (qPCR) and ChIP-PCR

Total RNA was extracted from cells or tissues using TRIzol Reagent (Invitrogen). The RNA was then reversely transcribed to cDNA using Primescript RT reagent Kit Thermo. To evaluate the mRNA levels of a number of genes, qPCR was performed on a CFX96 q–PCR System using SYBR Green qPCR master mix (Promega, USA). GAPDH was used as the internal control. All of the samples were normalized to internal controls, and fold changes were calculated based on relative quantification (2−△△Ct). Sequences of Primers were listed in **Supplementary Table 3**.

ChIP-PCR was performed using the Pierce Agarose Thermo ChIP Kit (Thermo) according to the manufacturer’s instruction. Briefly, cross-linking was performed by adding formaldehyde (final concentration 1%) and incubated at room temperature for 10 mins. Cross-linking reaction was terminated by the addition of glycine solution. Cells were washed with ice-cold PBS containing 0.1 mM PMSF. Cell pellets were collected by centrifugation at 3000g for 5 mins and resuspended in 1 ml of ChIP sonication buffer. DNA was sheared by sonication and the cell debris was pelleted by centrifugation at 9,000 g for 3 minutes. Equal aliquots of chromatin supernatants were subjected to overnight immunoprecipitation with anti-TRα antibody (Abcam) or IgG antibody (negative control). The primer sets used for PCR were listed in **Supplementary Table 3**.

### MRI

MRI experiments were performed on a 3-Tesla MRS*DRYMAG (Guildford, UK) preclinical scanner with a 17-cm-wide bore and a 20-mm birdcage mouse head coil. A T2-weighted two-dimensional fast spin-echo sequence was used for scanning mouse brain with the following parameters: scan direction = sagittal; repetition time = 5,000 ms; effective echo time (TEeff)/base echo time (TEbase) = 68 ms/17 ms; echo train length = 8; number of averages = 3; field of view = 25mm × 25 mm; matrix size = 256 × 240; slice thickness = 0.5 mm; gap = 0 mm; number of slices = 24. The tumor volume was assessed in each MRI examination independently by ROI-based volumetry. For the ROI-based measurement, Digital Imaging and Communications in Medicine (DICOM) imported into ImageJ. The areas which are traced on each sagittal T2-weight image are summed to calculate tumor volume (V). The tumor area (A) in each MR slice, 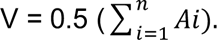 n, number of slices; *i*, individual slice number.

### Intracranial transplantation, in-vivo drug treatment and bioluminescent imaging (BLI)

Tumor cells were injected into the cerebella of *CB17/SCID* mice using a stereotaxic frame with a mouse adaptor (David Kopf Instruments), as described previously ^59^. 2 X10^5^ tumor cells suspended in 5µl NB-B22, were injected into each recipient cerebellum. Survival was defined as the days elapsed from transplantation until symptom onset.

In vivo T3 treatment was performed by I.P. injection (200ng/g, twice a day), which was initiated after tumors were established in Math1-Cre/Ptch1^loxp/loxp^ mice (4 weeks of age), Math1-Cre/LSL-SmoM2 mice (2 weeks of age), PDXs (luminescence signals in the range of 105–10^6^ rad/s). Equal volume of PBS was also injected as a control. Math1-Cre/LSL-SmoM2 mice were also treated with 50mg/kg vismodegib (twice a day) by I.P. injection. Body weight was routinely monitored for evaluation of possible T3 toxicities. T3 levels in the plasma and tumor tissues were measured at 2 hrs following T3 or PBS treatment, by using a T3 enzyme-linked immunosorbent assay kit (#EIAT3C, ThermoFisher). The blood was extracted from mice after the treatment, which was centrifuged at 3000 g for 15 mins to harvest the plasma after removal of cells and cell debris. Tumor tissues were homogenized in RIPA buffer and centrifuged at 3000 g for 15 mins, to remove the tissue debris. T3 levels were measured using the above kit according to the manufacturer’s instructions.

For BLI, tumor-bearing mice were injected by I.P. with 10 μl/g (body weight) of luciferin. BLI was performed at 10 min (efflux) after substrate injection using the IVIS Lumina III imaging system. Grayscale photographic images and bioluminescent color images were superimposed using LIVINGIMAGE (version 2.12, PerkinElmer) and IGOR Image Analysis FX software (WaveMetrics, Lake Oswego, OR). BLI signals were expressed in units of photons per cm^2^ per second per steradian (P/cm2/s/sr). All mice were anesthetized using 1–2% isoflurane gas during imaging.

### ECG recording

ECGs were recorded in mice using an ECG recording platform (Mouse Specifics, Boston, MA). The size and arrangement of the electrodes were configured to contact three paws, providing an ECG signal equivalent to Eithenoven lead I. To minimize the stress, mice were accustomed to the platform by placing them on it for at least 10 mins before ECGs were recorded. ECG signals were digitized and only data from continuous recordings of 20–30 signals were used in the following analyses. Each signal was analyzed using e-MOUSE (Mouse Specifics). The QRS duration, P-R interval, and heart rate (HR) were measured and reported automatically. To eliminate circadian influences, ECGs were recorded between 10:00 AM and 11:00 AM.

### Toxicity studies of T3 in tumor-bearing mice

***Math1-Cre/Ptch1^loxp/loxp^*** mice (3 weeks of age) were administered PBS or T3 at doses of 2 ng/g, 20 ng/g, or 200 ng/g or 2 μg/g (body weight) via I.P. injection, once daily, for consecutive 3 weeks. To minimize the circadian influences, mice were treated with T3 between 11:00 AM and 12:00 PM. To eliminate sex difference, only female mice were used in the toxicity studies.

Body weight, ECG recording, food consumption and water consumption were measured every XX days during the treatment. Blood samples and tumor tissues were collected at 24 hrs following the final T3/PBS treatment, for examining T3 levels in the plasma and tumor tissues, by using a T3 enzyme-linked immunosorbent assay kit (#EIAT3C, ThermoFisher), or analyzing tumor cell differentiation by immunohistochemistry. For T3 measurement, blood sample was centrifuged at 3000 g for 15 mins to harvest the plasma after removal of cells and cell debris. Tumor tissues were homogenized in RIPA buffer and centrifuged at 3000 g for 15 mins, to remove the tissue debris. T3 levels were measured using the above kit according to the manufacturer’s instructions.

Blood samples for hematological analyses were collected in Microtainer Blood Collection Tubes with K2EDTA (BD, USA). Peripheral blood was analyzed with a VetScan HM5 (Abaxis) hematology analyzer and discrimination between cell types was achieved by size. The following parameters were analyzed: lymphocytes, monocytes, and neutrophils, all reported as absolute cell counts as well as percentage of total white blood cells, red blood cells, hemoglobin, hematocrit, mean corpuscular volume (MCV), mean corpuscular hemoglobin (MCH), mean corpuscular hemoglobin concentration (MCHC), and red blood cell distribution width (RDWc), platelets (PLT), plateletcrit (PCT), mean platelet volume (MPV), and platelet distribution width (PDWc).

Major organs including brain, thyroid, lung, heart, liver, spleen, kidney and GI track were harvested in mice after the completion of T3/PBS treatment for 3 weeks. Histological slides prepared from these organs, were independently analyzed by two pathologists.

### RNA sequencing

Total RNA was extracted from tumor cells treated with T3 or PBS. Strand-specific mRNA-seq libraries for the Illumina platform were generated and sequenced following the manufacturer’s protocol. High-quality total RNA was used as input for the so-called dUTP library preparation method. Briefly, the mRNA fraction was purified from total RNA by polyA capture, fragmented and subjected to first-strand cDNA synthesis with random hexamers in the presence of Actinomycin D. The second-strand synthesis was performed incorporating dUTP instead of dTTP. Barcoded DNA adapters were ligated to both ends of the double-stranded cDNA and subjected to PCR amplification. The resultant library was checked on a Bioanalyzer (Agilent) and quantified. The libraries were multiplexed, clustered, and sequenced on an Illumina NextSeq 2000. The sequencing run was analyzed with the Illumina BCL-Convert, with demultiplexing based on sample-specific barcodes. The raw sequencing data produced was processed after removing the sequence reads which were of too low quality and discarding reads containing adaptor sequences. Raw data of RNA sequencing in this study are deposited in GEO under accession number: GSE224974.

### Statistical analysis

Unless stated otherwise, Student’s *t* test or one-way ANOVA was performed to determine the statistical significance of the difference. p<0.05 was considered statistically significant (*, *p*< 0.05; **, *p*< 0.01; ***, *p*< 0.001; ns, not significant). Error bars represent the SEM. Overall survival was assessed using the Kaplan-Meier survival analysis and the Mantel-Cox log-rank test was used to assess the significance of difference between survival curves. Data handling and statistical processing was performed using Graphpad Prism Software.

## Acknowledgement

We appreciate Drs. E. Golemis, J. Whestine and R. Locke for their insightful discussions and comments on the concepts and methodology of the work. We thank Drs. S. Liao for RNA sequencing analysis; J. Oesterling for flow cytometric analysis; A. Efimov for microscopy analysis; Y. Yang and D. Cvetkovic for MRI analysis. This research was supported by funds from National Cancer Institute (R01CA276273, ZJY), the CURE grant from Pennsylvania Department of Health (#4100085739, ZJY), American Cancer Society (ACSDG1900025, ZJY), the Andrew McDonough B+ Foundation (#1027381, ZJY), and the Buck County Board of Association at Fox Chase Cancer Center (ZJY), National Key Research and Development Program in China (2022YFE0133400, QL), National Natural Science Foundation of China (82172834, 82203397, QL) and the Core Comprehensive Cancer Center Grant CA06927 in support to Fox Chase Cancer Center’s facilities including Light Microscopy, Histopathology, Cell Culture, Biostatistics and Bioinformatics and the Talbot Library. Some graphics were created with BioRender (www.BioRender.com).

## Author Contributions

Z.Y. and Y.Y. conceived the project. Z.Y. and Y.Y. conceptualized and designed the majority of the experiments. Y.Y., S.A.V.-R., Q.L., Y.L., J.T. and Y.T. performed the experiments and interpreted the data. T.C. and R.J.W.-R. provided critical reagents. C.A.K. and S.C. contributed to analyze and interpret thyroid hormone measurement. Y.R., S.W. and K.C. conducted the histological analyses. S.R.H conducted the cardiac electrophysiology analyses. Z.Y. and Y.Y. wrote the manuscript and designed the figures.

## Figure Legends

**Supplementary Figure 1.**
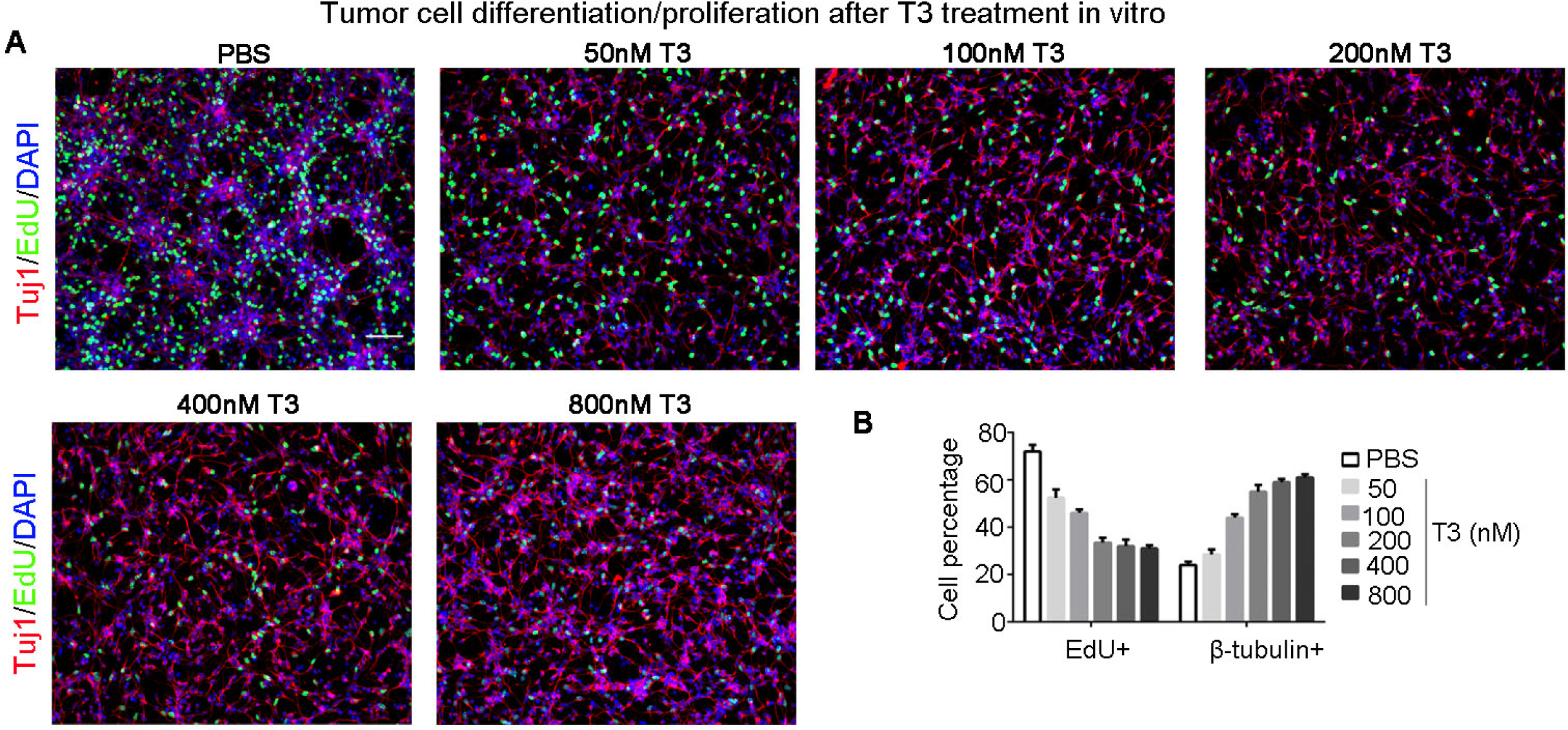
Terminal differentiation of MB tumor cells after T3 treatment. Tumor cells isolated from ***Math1-Cre/Ptch1^loxp/loxp^*** mice, were treated with T3 at designated concentrations, for 48 hrs. After being pulsed with EdU for 2 hrs, tumor cells were harvested to examine cell proliferation (EdU incorporation) and differentiation (β-tubulin expression) by immunocytochemistry (**A**). DAPI was used to counterstain cell nuclei. The percentage of EdU+ cells or β-tubulin+ cells was quantified (**B**). Scale bar: 50µm (**A**)

**Supplementary Figure 2.**
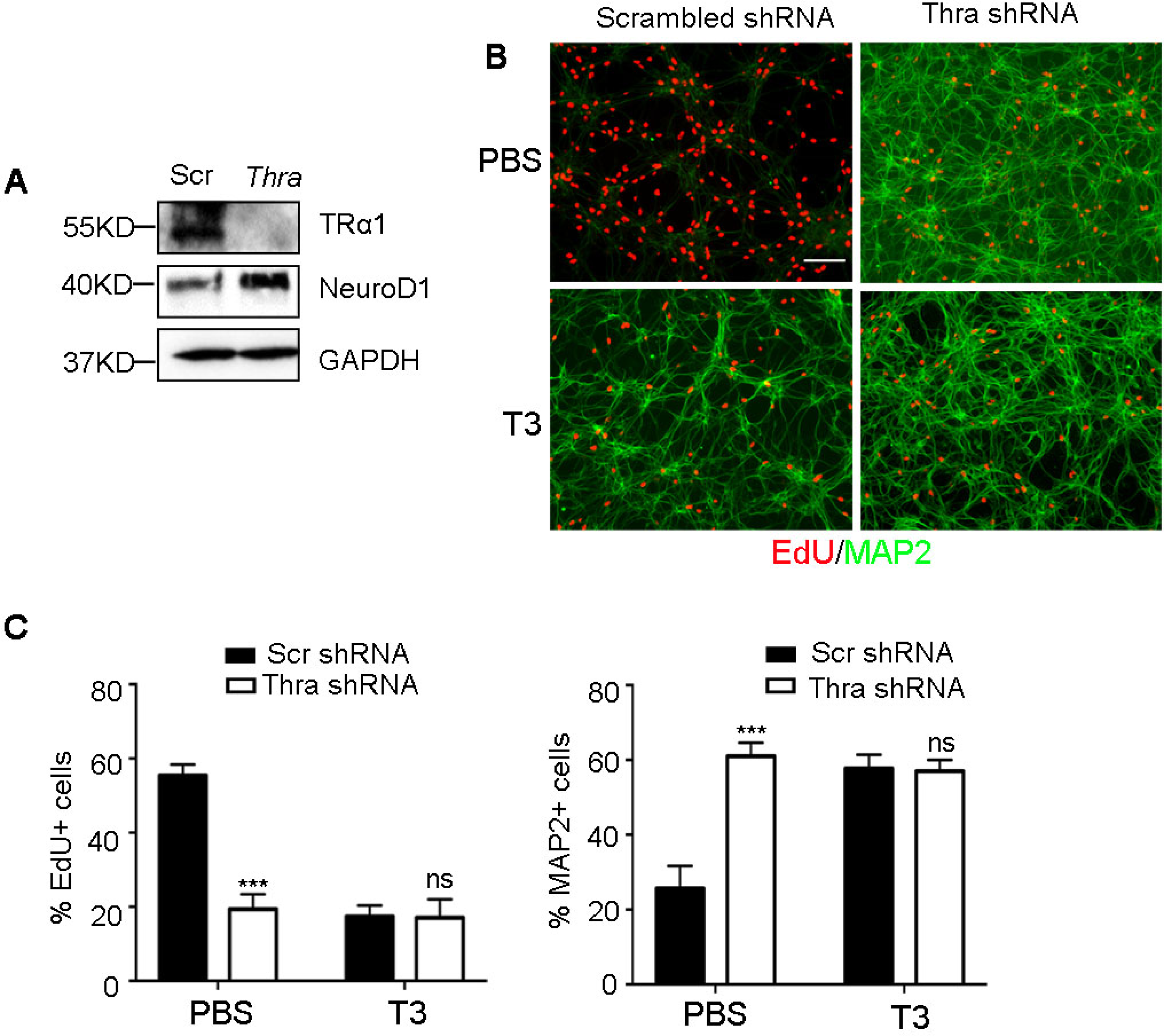
*Thra* knockdown induces tumor cell differentiation. Tumor cells were infected with a lentivirus carrying a shRNA specific for *Thra*, or a scrambled shRNA, and harvested at 48 hrs following the infection, to examine protein levels of TRα1, NeuroD1 and GAPDH by western blotting (**A**). Before being harvested, tumor cells were pulsed with EdU for 2 hrs. Tumor cell proliferation (EdU incorporation) and differentiation (MAP2 expression) were examined by immunocytochemistry (**B**). The percentage of EdU+ cells or MAP2+ cells in tumor cells transduced with *Thra* shRNA or scrambled shRNA, was quantified (**C**). Scale bar: 50µm.

**Supplementary Figure 3.**
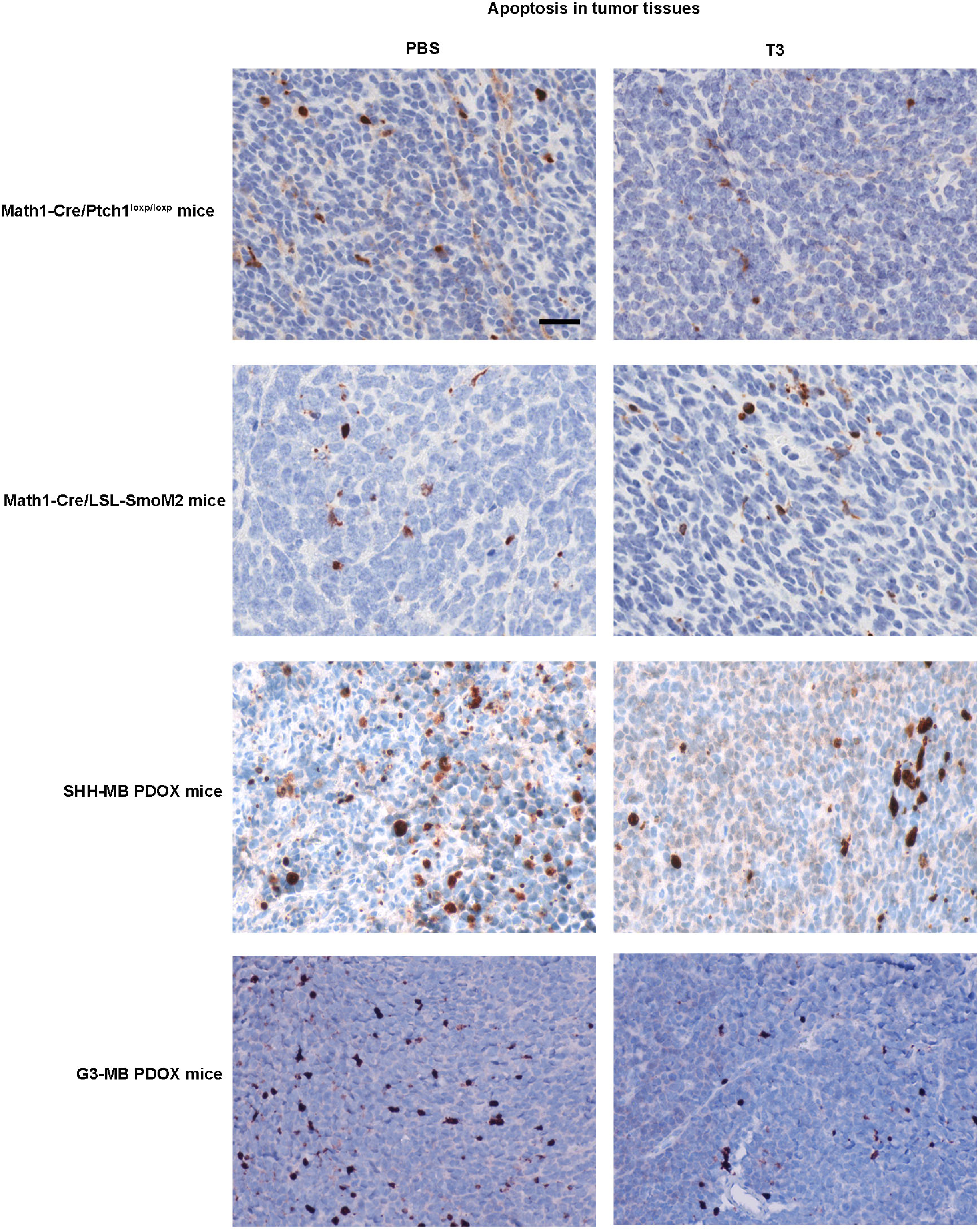
No increased apoptosis in MB tissues after T3 treatment. MB tissues were harvested from ***Math1-Cre/Ptch1^loxp/loxp^*** mice, ***Math1-Cre/LSL-SmoM2*** mice, SHH-MB PDOX-bearing mice and G3-MB PDOX-bearing mice after treatment with T3 or PBS. Tumor sections were immunostained with an antibody against cleaved caspase-3. Scale bar: 100µm

**Supplementary Figure 4.**
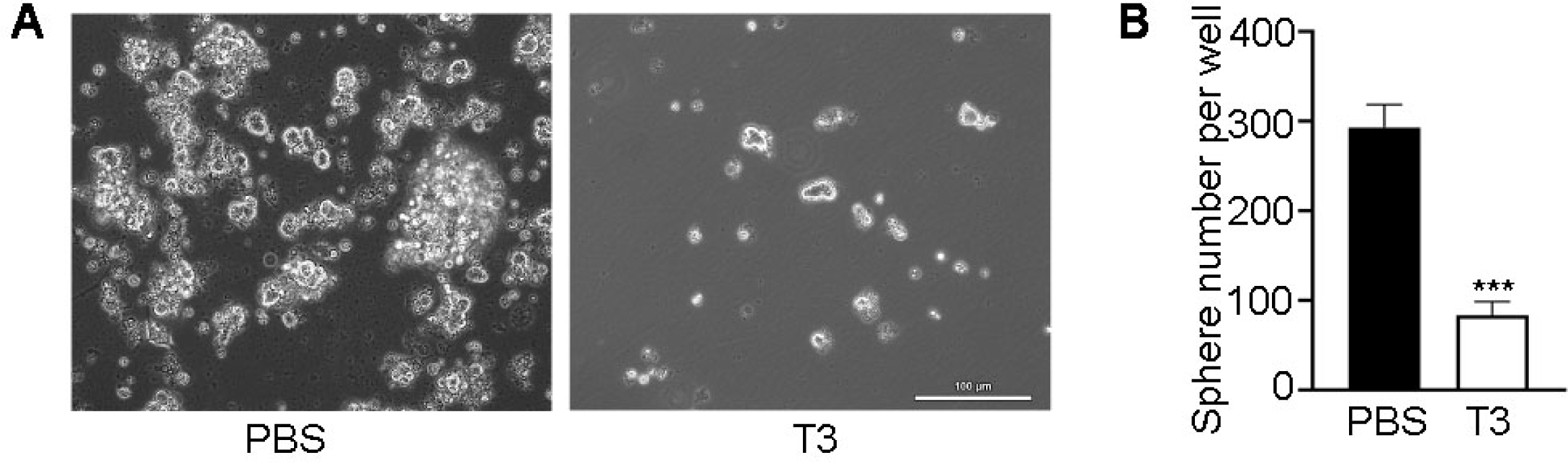
Tumoroid formation of RCMB28 cells after treatment with T3 or PBS. RCMB28 cells were cultured under sphere-forming condition for 5 days, in the presence of T3 or PBS (**A**). Number of tumoroids was counted under a microscope (**B**). Scale bars: 100µm

**Supplementary Videos 1-2**

Littermates of *Math1-Cre/Ptch1^loxp/loxp^* mice were treated with T3 or PBS, for 2 weeks. The movement of treated mice was recorded.

**Supplementary Videos 3-5**

Littermates of *Math1-Cre/LSL-SmoM2* mice were treated with T3, PBS or vismodegib for 2 weeks. The movement of treated mice was recorded.

**Supplementary Videos 6-10**

Littermates of *Math1-Cre/Ptch1^loxp/loxp^* mice were treated with PBS or T3 (2ng/g, 20ng/g, 200ng/g, 2µg/g) for 3 weeks. The movement of treated mice were recorded.

**Supplementary Table 1**

Transcripts differentially expressed between T3-treated tumor cells and PBS-treated tumor cells.

**Supplementary Table 2**

Hematology of mice after the treatment with T3/PBS for 3 weeks

**Supplementary Table 3**

Sequences of PCR primers, shRNAs and gRNAs used in this study.

